# VCAb: A web-tool for structure-guided antibody engineering

**DOI:** 10.1101/2024.06.05.597540

**Authors:** Dongjun Guo, Joseph Chi-Fung Ng, Deborah K. Dunn-Walters, Franca Fraternali

**Author notes:** Shared first authorship (Dongjun Guo, Joseph Chi-Fung Ng).

## Abstract

Effective responses against different immune challenges require secretion of antibodies with various isotypes performing specific effector functions. Structural information on these isotypes is essential to engineer antibodies with desired physico-chemical features of their antigen-binding properties, and optimal stability and developability as potential therapeutic antibodies. *In silico* mutational scanning profiles on antibody structures would further pinpoint candidate mutations for enhancing antibody stability and function. Although a number of antibody structure databases exist, a public data resource which provides clear, consistent annotation of isotypes, species coverage of 3D antibody structures and their deep mutation profiles is currently lacking. The <u>V</u> and <u>C</u> region bearing <u>a</u>nti<u>b</u>ody (VCAb) web tool is established with the purpose to clarify these annotations and provide an accessible and easily consultable resource to facilitate antibody engineering. VCAb currently provides data on 6,948 experimentally determined antibody structures including both V and C regions from different species. Additionally, VCAb provides annotations of species and isotypes with both V and C region numbering schemes applied, which can be interactively queried or downloaded in batch. Multiple *in silico* mutational scanning methods are applied on VCAb structures to provide an easily accessible interface for querying the impact of mutations on antibody stability. These features are implemented in a R shiny application to enable interactive data interrogation. VCAb is freely accessible at https://fraternalilab.cs.ucl.ac.uk/VCAb/. The source code to generate the VCAb database and the online R shiny application is available at https://github.com/Fraternalilab/VCAb, enabling users to set up local VCAb instances.

## Introduction

Antibodies, a key component of the immune system, are composed of two pairs of heavy (H) chain and light (L) chain, with each chain bearing variable (V) and constant (C) regions (Dreyer and Bennett, 1965; Lu et al., 2018; Chiu et al., 2019; Guo et al., 2024). The V region directly interacts with the antigen, while the C region of H chain, which defines the “isotype” of the antibody, determines its relevance in different immune processes. Antibodies are broadly applied as therapeutics because of its specificity to the targeted antigens and the effector functions it triggers to coordinate immune clearance of such antigens. A major research focus in the field of antibody engineering has been to evolve the binding affinity to antigen by changing the V region, due to its direct role in engaging the antigen (Tabasinezhad *et al*., 2019; Hie et al., 2023). Unstable antibodies displayed impeded or lost efficacy, high chances of aggregation, low production yield, and low propensity in becoming developable therapeutic antibodies (Ma *et al*., 2020; Hanning *et al*., 2022). To this end, biophysical, energy-based approaches and machine-learning based methods have been used to generate large-scale *in silico* predictions of mutations to improve antibody thermostability (Leaver-Fay *et al*., 2011; Ruffolo et al., 2021; Cheng et al., 2023; Harmalkar et al., 2023). However, studies have highlighted the importance of the C region in modulating antigen interactions (Cooper *et al*., 1993; Torres et al., 2007; Tudor *et al*., 2012; Lua et al., 2018; Khamassi et al., 2020; Casadevall and Janda, 2012; Guo et al., 2024) and fulfilling antibody stability and function, underscoring the necessity to consider entire antibody structures during antibody engineering. In addition, the development of next-generation sequencing methods has allowed for deep sampling of the antibody repertoire of many individuals, profiling a great diversity of V regions coupled with different isotypes (Marks and Deane, 2020; Olsen et al., 2022). This raises questions on the properties of these antibodies at the protein structural level, and the scope to engineer both V and C regions to improve the binding functions and stability of the antibody. To answer these questions, collection and annotation of antibody structures containing both V and C region, with different types of H chain and L chain, are needed.

A number of databases are available for interrogating antibody structural data, such as Protein Data Bank (PDB) (Berman *et al*., 2003), IMGT (ImMunoGeneTics information system)/ 3Dstructure-DB (Kaas *et al*., 2004; Ehrenmann and Lefranc, 2011) and SAbDab (Dunbar *et al*., 2014; Schneider et al., 2022), amongst others. Whilst the PDB encompasses the breadth of protein structures, it lacks an easy-to-use query interface for the user to specifically filter antibody structures. On the other hand, other well-established databases like IMGT/3Dstructure-DB do not offer a programmatic interface to access data in bulk, limiting queries to one structure at a time, precluding large-scale analyses pipeline integration. A recently antibody-based queriable database, SAbDab (Dunbar *et al*., 2014; Schneider et al., 2022), was specifically built to offer accurate annotation of V regions of antibody structures. To the best of our knowledge, currently available databases do not provide information of the effect of mutations in modulating stability. To address the gap for reliable, large-scale and easily retrievable annotations of both V and C region structures and their *in silico* mutational scanning profiles, we present VCAb (<u>V</u> and <u>C</u> region bearing <u>a</u>nti<u>b</u>ody database). VCAb collates experimentally resolved antibody structures with both V and C regions; the database (a) is readily updated and easily queried, (b) contains clear information about sequence, isotype/light chain type and structural coverage, (c) offers, in addition to V region sequences conforming to the IMGT numbering scheme provided by other standard tools (Abhinandan and Martin, 2008; Dunbar and Deane, 2016a; Giudicelli *et al*., 2011), IMGT-gapped C region sequences, allowing consistent analysis of structural features such as domain packing geometries, and uniquely, (d) *in silico* mutational scanning data for experimental antibody structures which would be of useful help in designing stable antibodies. VCAb can be queried online (https://fraternalilab.cs.ucl.ac.uk/VCAb/) using characteristics such as isotype, sequence similarity, or CH1-CL interface similarity. VCAb can serve as an ideal tool for researchers interested in the selection of template for antibody modelling purposes, and the structural properties of the antibody to optimise the geometries and stabilities of antibody designs (**Figure 1a**).

**Fig. 1:**
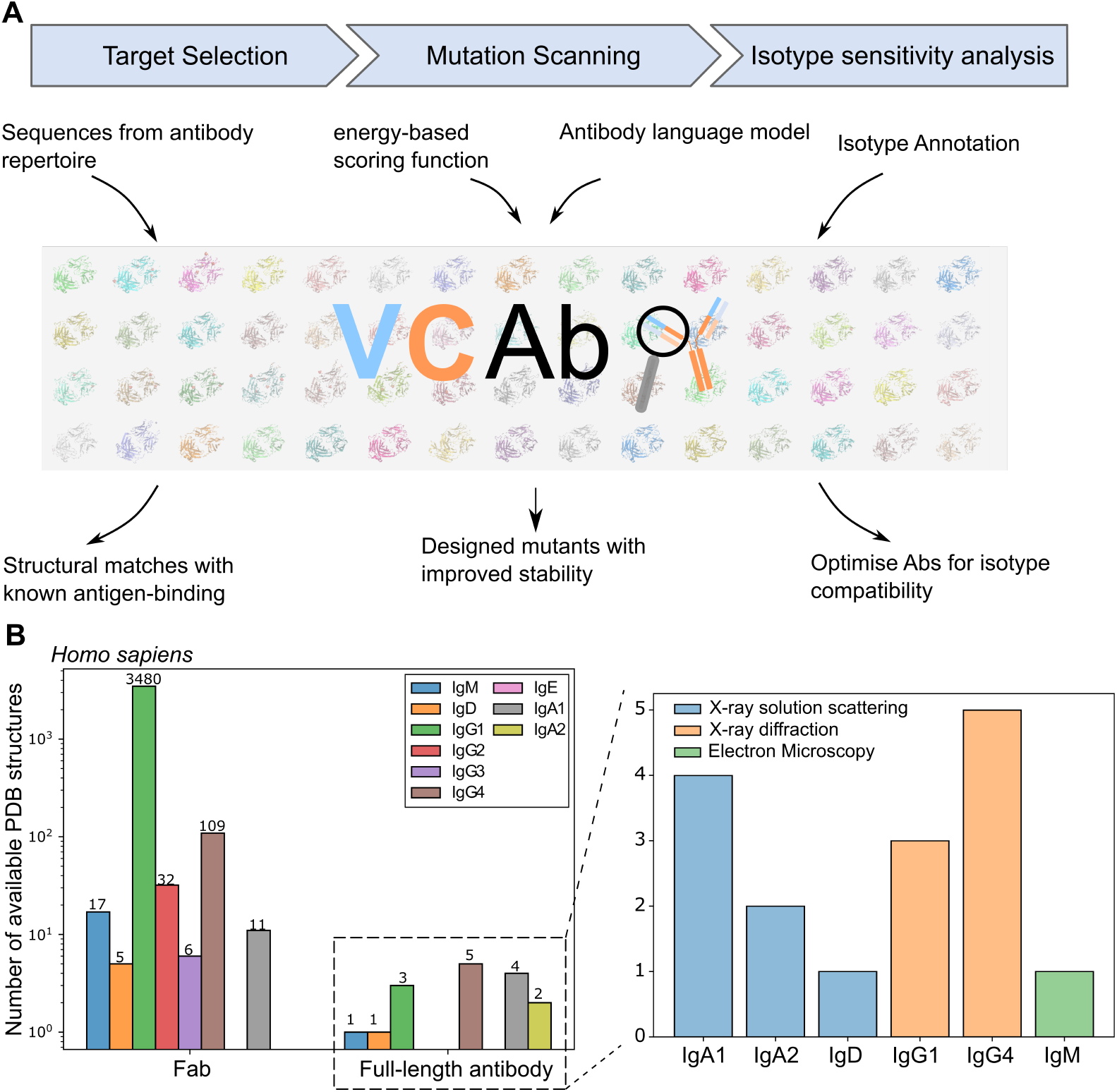
VCAb: an user-friendly web tool to guide antibody engineering. (A) VCAb offers functionalities to facilitate multiple steps involved in antibody engineering, including target selection, mutational scanning and isotype sensitivity analysis. (B) Experimental human antibody structures of different isotypes, structural coverages are collected in VCAb. The bar charts show (*left*) statistics on the coverage of human antibody structures and (*right*) the isotype distribution of full-length human antibody structures, as at April 29th, 2024.

## Implementation

### Data Collection

SEQRES records of protein chains with resolved structures were automatically downloaded from worldwide PDB archive (Burley *et al*., 2019). Antibody sequences with both V and C regions were identified from the downloaded protein sequences using ANARCIvc (Supplementary S3.1.3), a package we modified from ANARCI (Dunbar and Deane, 2016b) to number sequences of both V and C regions conforming to IMGT rules. Sequences successfully analysed and numbered using ANARCIvc were deemed to contain the correct sequence features expected for antibody V and C regions. Downloaded mmCIF antibody structure files were annotated with structural metadata using the PDBe application programming interface (API); all procedures were automated in a set of python scripts available in the VCAb github repository. As of 29th April 2024, there were 6,948 antibodies collected in VCAb, 3,676 of which are human (Downloaded on Apr.29th, 2024, **Figure 1b**). This data collection process has been automated to update the database monthly.

### Features annotation

VCAb annotates antibody species, isotype, and structural coverage by comparing its sequence to all IMGT (Lefranc *et al*., 2015) reference alleles using BLAST (Altschul *et al*., 1990a). To address spurious species annotation provided by the PDBe API whilst mitigating potential artefacts arisen from BLAST local alignments, we considered each V/C domain separately, consolidated both annotations and overwrote the PDBe-provided species if the best BLAST hit has a larger percentage identity by 8% compared to the PDBe annotation; we observed that this cutoff effectively separated antibody V and C sequences from different species (Supplementary S3.1.2). We further consolidated the nominated species for each domain to generate a final annotation which flagged different engineered formats (humanised, chimera). Isotype and light chain type were identified to be the BLAST hit with highest percentage identity. To ensure the accurate assignment of structural coverage, the sequence representing residues with ATOM records extracted from mmCIF was used as input to BLAST alignments against IMGT reference alleles. We classified structures as Fab or full antibody, depending on whether coordinates for CH2, CH3, or CH4 were included in the structure. We provided the following annotations of Fab structures: (1) packing angles (elbow angle, CH1-CL angle) as defined by Fernández-Quintero *et al*. (2020); (2) annotations of interface between heavy and light chains (“H-L interface”) derived using POPSCOMP (Kleinjung and Fraternali, 2005); (3) contact matrix considering the C*α*-C*α* distances between all residues along the heavy and light chains. All CH1 and CL sequences were numbered using ANARCIvc (Supplementary S3.1.3) so that all downstream analyses conform to the IMGT standard numbering scheme for the C region for ease of comparison.

### In silico antibody mutational scanning

We applied three *in silico* mutational scanning methods to evaluate mutations which can affect the stability of the antibody: this includes (1) Rosetta point mutant scan application (Leaver-Fay *et al*., 2011) yielding structure-based scores focusing on the physico-chemical features of residues; (2) pseudo-log-likelihood from the AntiBERTy (Ruffolo *et al*., 2021), a language model for the antibody V region of antibody. We scaled the raw AntiBERTy score of every point mutation by subtracting the AntiBERTy score for the wild type (WT) residue at each position, such that they can be interpreted as relative changes in amino acid preference compared to WT. This method has been applied to predict mutational effects on protein function (Meier *et al*., 2021). (3) AlphaMissense (Cheng *et al*., 2023) scores to evaluate mutational effects of C-region mutations. These methods were applied on any possible amino acid substitution in every VCAb entry to constitute *in silico* mutational scanning datasets, with all of them freely accessible for download.

### VCAb web server

The VCAb website has been built to allow data access for academic research purposes, available at https://fraternalilab.cs.ucl.ac.uk/VCAb/. The website displays the features described above for each VCAb entry, enables filtering of antibody structures based on these features, and supports searches by sequence similarity, using BLAST (Altschul *et al*., 1990b) accessed via the rBLAST package (version 0.99.2). This search is flexible to the region of interest (V region or both V & C regions) and both paired and unpaired H/L chain sequences, and supports input of single sequences, uploads of multiple FASTA sequences (maximum 200 per batch), as well as tabulated antibody repertoire data in standard formats (AIRR standard tab-separated files, comma-separated files, and output from single-cell repertoire sequencing analysis generated by the 10x Genomics Cellranger software). Users can also search VCAb by CH1-CL interface similarity, which is derived by comparing the contact matrix at the CH1-CL interface generated for each VCAb entry (Supplementary Materials section S1.2.2).

The VCAb webserver provides visualisation functionalities for 3D structures, structural coverage, antibody numbering information for both V and C regions, as well as tabulated details of H-L interface, disulphide bonds and *in silico* mutational scanning results. The interactive 3D viewer enables detailed inspection of the structure. The following metadata are displayed for each VCAb entry: PDB and chain identifiers, assigned germline CH and CL alleles, structural coverage, species. A list of detailed information included in VCAb can be found in the online documentation accessible in the ‘About’ page on the online interface; these additional columns can be accessed either by customising the table view to show these additional columns, or by bulk downloads of search results and/or the entire VCAb database in Comma Separated Value (CSV) format.

## Application

### Investigation of COVID-19 repertoire to illustrate the binding between antibody and RBD domain

The availability of antigen-antibody complex structures in VCAb allows for annotation of the likely antigen-binding mechanisms for sequences without experimental investigations of their binding properties (**Figure 2A**), for example those obtained via high-throughput sequencing of the antibody repertoire. The volume of such data is rapidly expanding, yielding paired heavy and light chain immunoglobulins attributed to single B cells (Olsen *et al*., 2022) collected from vaccinations and disease scenarios (Schultheiß *et al*., 2020; Kotagiri et al., 2022; Jin et al., 2021; Stewart et al., 2022). VCAb can help here in annotating these sequences to their closest structural match, thereby closing the sequence-structure gap and potentially assisting interpretation of their functional properties (e.g. antibody-antigen interaction). We used repertoire data from Stewart et al. (2022) of hospitalised Coronavirus disease 2019 (COVID-19) patients, and applied VCAb to search for structural matches and illustrate how the sampled antibody sequences bind to the antigen, the spike protein S1 of severe acute respiratory syndrome coronavirus 2 (SARS-CoV-2). The top four structural hits for the sampled sequence show high sequence similarity (larger than 94%) for the V region, with all of them containing the antigen (spike protein S1), which indicates the likely antigen for this selected antibody sequence from the repertoire (**Figure S2A**). Another sampled sequence gives rise to multiple structural hits with the S1 protein as well, suggesting the identity of the antigen for the selected sequence entry (**Figure S2B**).

**Fig. 2:**
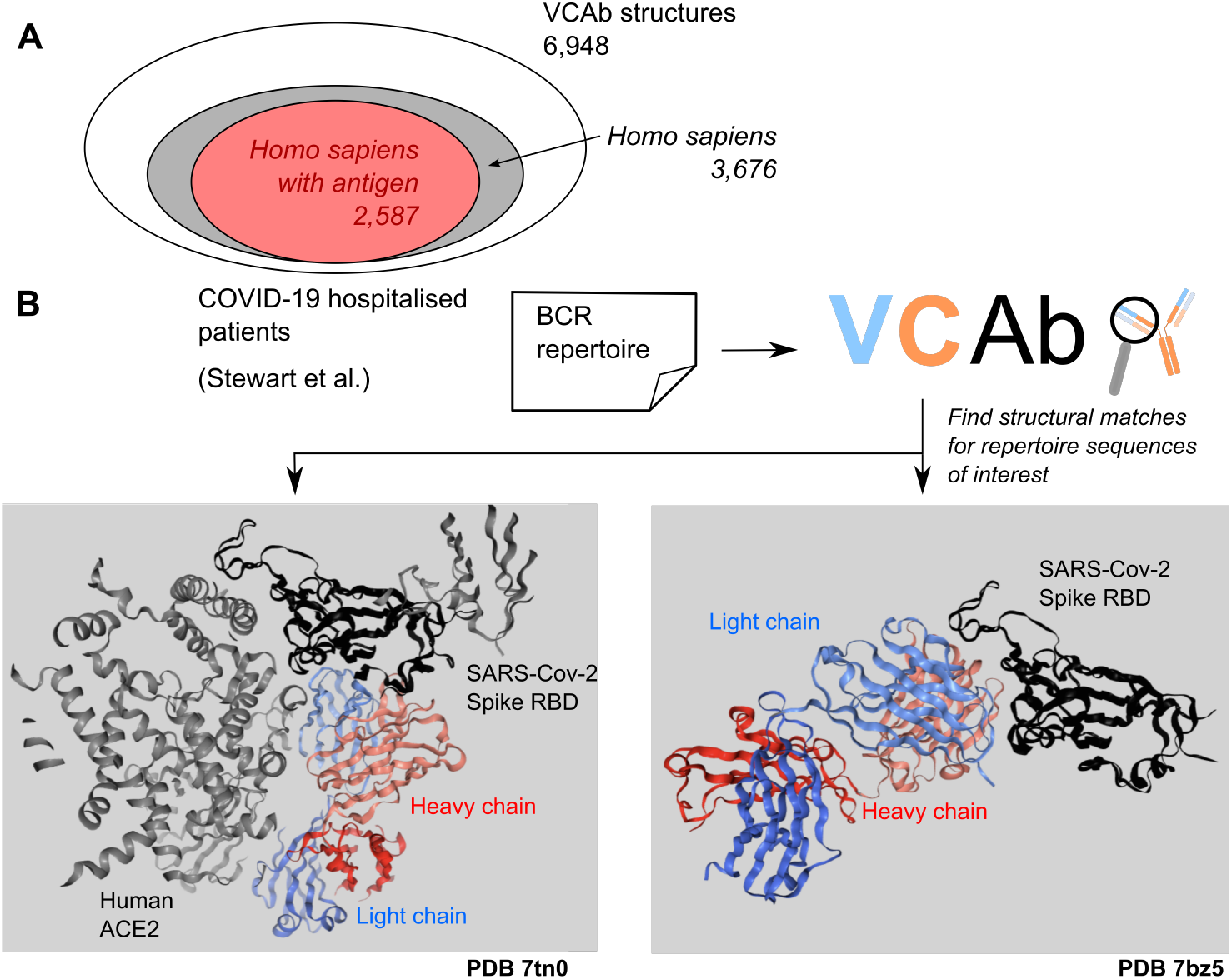
Investigation of COVID-19 repertoire from a structural perspective. **(A)** Antibody structural space in VCAb. The majority (52.9%) of antibody structures in VCAb is from human, among which 2,587/3,676 (70.4%) of these are co-complexed with antigens. **(B)** Two sequences from the sampled repertoire (Stewart *et al*., 2022) are inputted into VCAb to search for matches in the dataset of resolved antibody structures. We obtained a structural match (*left*, PDB 7tn0) showing the recognition of the RBD epitope distinct from ACE2 binding site. For the other sequence (*right*), the corresponding structure match (PDB 7bz5) shows this antibody directly blocking ACE2 binding.

When we used the structural viewer functionality provided in VCAb to compare the structure matches ranked first (7bz5 HL and 7tn0 MN) for these two selected repertoire sequences, it comes to our attention that they interact with the same antigen differently. 7bz5 HL binds to the cellular receptor ACE2 (Angiotensin-converting enzyme 2) binding site of the spike receptor binding domain (RBD), making contact with most residues on the RBD-ACE2 interface for its competition with ACE2 (Wu *et al*., 2020). On the other hand, 7tn0 MN binds to the cryptic epitope at the other side of the RBD domain without occupying the ACE2 binding site (McCallum *et al*., 2022) (**Figure 2AB**). Here, VCAb allows the user to interpret the epitope for these two antibodies and their likely modes to efficiently engage the spike protein (Piccoli *et al*., 2020).

### *In silico* mutational scanning of Antibody structures

Thermostability is one of the parameters used as a guidance in therapeutic antibody design to achieve suitable candidates for further development (Hanning *et al*., 2022). *In vitro* deep mutational scanning to measure the (thermostability) impact of point mutations is a costly and time-consuming pipeline involving protein display, screening and sequencing, etc. (Hanning *et al*., 2022). Here, VCAb offers *in silico* predictions of single point mutations for each antibody structure generated using different methods: an energy-based method relying on the antibody structure (Rosetta, Leaver-Fay *et al*. (2011)), and machine-leaning based methods (using the antibody-specific language model AntiBERTy (Ruffolo *et al*., 2021) for V region and the AlphaMissense model (Cheng *et al*., 2023) for C region). This facilitates the prioritisation of promising mutations to be characterised experimentally.

MEDI8852 is an antibody neutralizing influenza A hemagglutinin (Hie *et al*., 2023) with an experimentally resolved structure (PDB 5jw5). **Figure 3A** shows the stability scores calculated from Rosetta and AntiBERTy, with red being unpreferable and blue being preferable. To validate the utility of these data, we compared these scores against a recent analysis (Hie *et al*., 2023) which used a protein language model (ESM-1b) to design novel mutations on its unmutated common ancestor (UCA) with high sequence similarity, and validated their impact experimentally. The mutation G95P in the VL domain was found to be destabilising (decrease in melting temperature [Tm]) but its affinity to the antigen was enhanced (Hie *et al*., 2023). Analysing this in VCAb, this mutation is predicted as destabilizing using both Rosetta and AntiBERTy, and is consistent with its experimentally validation with its ΔTm negative. In fact, most mutations at this position are predicted as unpreferable, with a few predicted as neutral by Rosetta. Glycine has the smallest side chain among all the amino-acids, and by inspecting it in the 3D structural viewer, it sits at the interface between VH-VL domain. This indicates the importance of the spatial localisation of this residue at the VH-VL interface and can act as the starting point for further analysis.

**Fig. 3:**
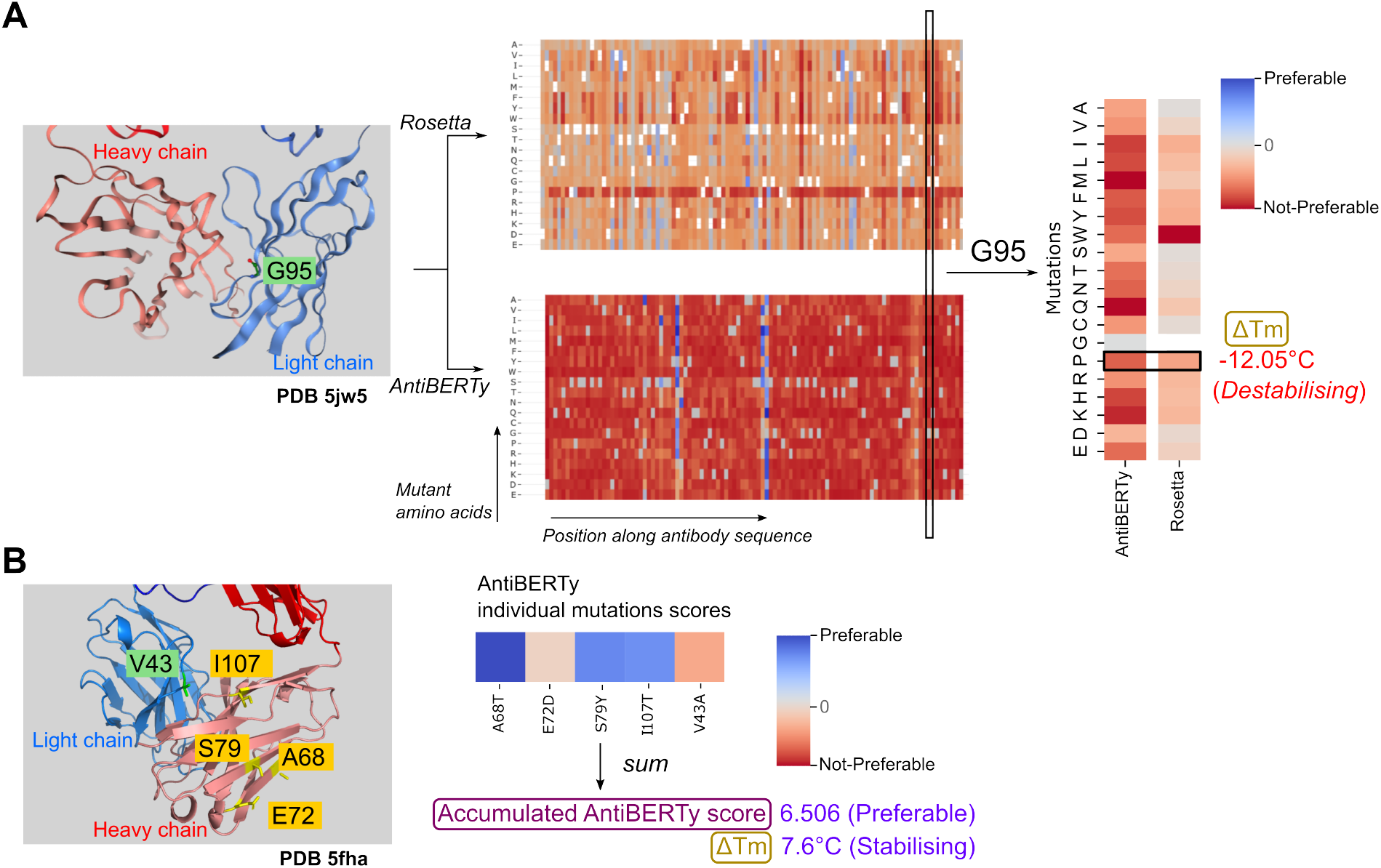
*In silico* mutational scanning of Antibody structures. (A) AntiBERTy and Rosetta scores for antibody (PDB 5jw5) at residue G95, with red being not-preferable and blue being preferable. Experimental measurement of the ΔTm for G95P shows a negative value (i.e. this mutation is destabilising to the antibody). The position G95 is highlighted (ball-and-stick in green) on the three-dimensional structure is shown on the left. (B) An example of antibody structure (PDB 5fha) with multiple mutations highlighted. Mutations on light chain is colored green, with mutations on heavy chain colored yellow. Pseudo-log likelihood for individual mutations are added together to estimate the effect of introducing multiple mutations together. Experimental measurements of this quintuple mutant show the changes of melting temperature of 7.6 ºC, indicating its stabilizing effect.

For each residue in the V region of any VCAb structure both Rosetta and AntiBERTY scores are provided to users for comparing the effects of using different types of information (sequence and structure) in predicting mutational impact. We note that for Rosetta, since each point mutation was predicted separately, the resultant scores on VCAb were not additive and therefore would not be suitable for predicting the effect of multiple mutations in combination. Here, AntiBERTY, being a language-model based method, can address this issue: since each amino acid is represented as a “word” in a sentence, the pseudo-log-likelihood returned by the model can be summed together to represent the likelihood of observing several amino acids in combination (Figure S4, Supplementary Materials), in the same way as the likelihood of a given sentence being presented is evaluated in language models used in natural language processing (NLP) (Meier *et al*., 2021; Salazar *et al*., 2020; Shin *et al*., 2019). mAb114 is an antibody binding to the glycoprotein of ebolavirus (PDB ID 5fha). A quintuple mutant (Heavy chain: A68T, E72D, S79Y, I113T; Light chain: V43A) has been designed with improved affinity to the antigen (Hie *et al*., 2023). Three out of the five single mutations are predicted as preferable by AntiBERty (Figure 3B). Summing over the pseudo-log likelihood of individual mutations yields a positive accumulated score for the co-occurrence of the five mutations together, indicating that it is preferable for the five mutations being presented at the same time, compared with the wild type. This prediction is confirmed by experimental measurement of thermostability indicated by positive ΔTm (**Figure 3B**).

### Exploring the consequence of isotype switching on antibody structural stability

Therapeutic antibody design requires careful selection of isotypes to achieve desired downstream effector functions. This is relevant also for *in vivo* antibody maturation, where isotype switching is a critical process to adapt the antibody to function in different contexts. However, how isotype switching would affect antibody stability has not been comprehensively analysed. Benefiting from the mutational scanning analysis and isotype annotation of the antibody structures in VCAb, user can start to investigate the hot-spots in V region which are sensitive (in terms of the changes of stability) upon coupling with different C regions, and potentially engineer these hot-spots to stabilise the antibody. Here we attempted isotype switching *in silico*, using a set of Fab structures in VCAb with identical V region (originally isolated from a lymphoma patient (Houdayer *et al*., 1993)) coupled with both IgA1 (PDB 3qnx) and IgG1 (PDB 3qo0) (**Figure 4A**). Comparing the Rosetta pmut scan results of the VH region, the method agrees on mutational impact at most positions in the two structures, although discrepancies exist for several mutations (**Figure 4B**). Using another IgG1 structure with the same V region (PDB ID 3qo1) as the negative control (**Figure 4B**), we considered the mutational impact upon switching isotypes to detect locations sensitive to the selection of isotypes (**Figure 4C**, see Supplementary Materials section S3.2). The fold difference between the IgA1-IgG1 comparison over the IgG1-IgG1 comparison is calculated to indicate the favorable isotype, with negative values meaning mutations at this position tends to be IgA1-favorable and positive values being IgG1-favorable. In **Figure 4D** we display VH residues with mutations showing significant differences in the scores calculated in the context of different isotypes. Most of these positions are located in the loops close to the C region, highlighting how V and C regions together determine structural stability. Interestingly, some residues on the CDRH3 loop are also highlighted from this analysis. Previous research discovered that antibodies with the same V region but different isotypes have distinct affinity to the antigen (Tudor *et al*., 2012; Casadevall and Janda, 2012). This analysis helps prioritising positions for further investigation in the relationship between antigen binding affinity and the sensitivity of these positions towards isotype switching.

**Fig. 4:**
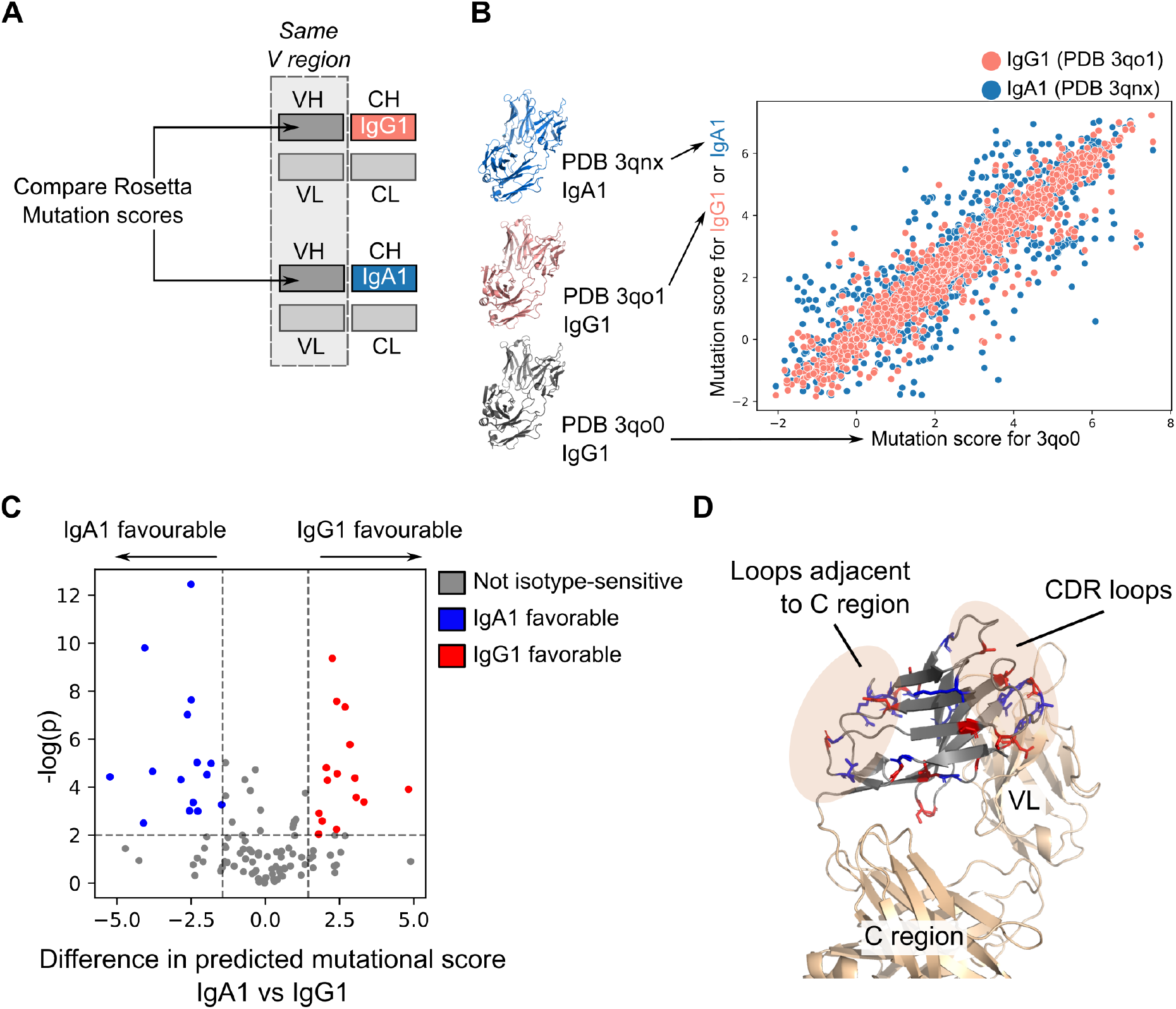
Exploring the consequence of isotype switching on antibody structural stability. **(A)** Rosetta mutational scanning scores for VH residues from antibodies of different isotypes are compared. These antibodies have the same VH and Light chain sequences, the only difference between them is their CH1 domain. **(B)** Scatter plot of Rosetta mutational scanning scores. Red dots correspond to the correlation between Rosetta mutational scores of structures of the same isotype (IgG1: PDB ID 3qo0 and PDB ID 3qo1), blue dots correspond to correlation between mutational scores calculated using structures of different isotypes (IgG1: PDB ID 3qo0; and IgA1: PDB ID 3qnx). **(C)** A volcano plot is derived for each position in the VH domain by comparing the difference in the mutation scores calculated from the IgA1 versus the IgG1 structure (see Supplementary Materials section S3.2). Blue data points indicate positions where mutations will favour IgA1 and red indicating positions where mutations will be IgG1 favorable. **(D)** The IgG1/IgA1-sensitive positions highlighted in panel (C) are visualised on the antibody structure. Most of them concentrate on CDR loops and loops close to C region.

## Conclusion

VCAb harmonises annotations of antibody isotypes, species and structural coverage, and provides data for detailed analysis of antibody stability. The VCAb database is updated once a month to include newly released, experimentally determined antibody structures. Users can interact with the online webserver for accessing annotations of individual structures, or for batch downloads of these annotations over many structures; alternatively, users can also use the publicly available source code to build and maintain a local version of the database for offline usage. We foresee that VCAb will provide useful information for antibody design and engineering, as well as the selection of homology modelling templates to investigate the structures of different antibody isotypes. The numbering of V and C regions enabled by ANARCIvc will allow further, large-scale mining of features at the V and C domains to inform antibody designs and understanding of the biophysical properties governing stability and function of antibody domains.

## Author contributions statement

DG collected data and generated the VCAb database and the shiny app, under the supervision of FF, DDW and JN. JN administered the shiny app on the webserver. DG and JN wrote the manuscript. All authors commented, revised and approved the manuscript.

## Data Availability

VCAb is freely accessible at https://fraternalilab.cs.ucl.ac.uk/VCAb/. The source code to generate the VCAb database and the online R shiny application is available at https://github.com/Fraternalilab/VCAb. The package ANARCIvc developed on the top of ANARCI is available at https://github.com/Fraternalilab/ANARCI_vc.

## Acknowledgments

We would like to thank all members of the Fraternali group for comments and suggestions. We also thank Jens Kleinjung for his advice on the shiny application. Rosetta point mutant application is run on the ARCHER2 UK National Supercomputing Service (https://www.archer2.ac.uk).

