## Supplementary Materials for "VCAb: A web-tool for structure-guided antibody engineering"

Fraternali^1,∗^

*^1^Institute of Structural and Molecular Biology, University College London, Darwin Building, Gower Street, London WC1E 6BT,United Kingdom*

*^2^Randall Centre for Cell and Molecular Biophysics, School of Basic and Medical Biosciences, King’s College London, London SE1 1UL, United Kingdom*

*^3^School of Biosciences and Medicine, University of Surrey, Guildford GU2 7XH, United Kingdom*

*^+^Shared first authorship.*

### S1 Description of functionalities

#### S1.1 Visualization of antibody information

Each Antibody entry in the VCAb containing features of four aspects: sequence, species, isotype, structure metadata, and three-dimensional structure.

User can customise the table view to include columns being displayed, with bulk downloads of search results and/or the entire VCAb database in Comma Separated Value (CSV) format enabled.

User can also query VCAb by filterring features of their interest. This could be the starting point of antibody engineering for the selection of potential structures as the template for further modification. Entries matching the user’s query is displayed in the antibody information panel. User can download these compiled data tables, visualise and inspect the selected antibody structure and interactively browse the CH1-CL interface residues in the 3D structure viewer, with all CH1-CL interface residues tabulated, Fab contact matrix displayed, and *in silico* mutational scanning profiles shown.

All the features listed below can be accessed via VCAb webserver. A list of detailed information included in VCAb can be found in the online documentation accessible in the ”About” page of VCAb.

##### S1.1.1 Sequence

The sequence for each antibody entry are provided and downloadable to the user. We also list the sequence represented in the PDB ATOM records. This reflects the true coverage in PDB structure, and is therefore relevant in structural investigations (e.g. in prioritising templates to be used to model antibody structures). Users can compare the domain coverage of the sequence downloaded from PDBe API and the sequence representing PDB ATOM records by clicking the “Sequence Coverage” tab.

Each sequence is numbered based on IMGT scheme (Lefranc *et al.*, 2005) by ANARCIvc package we developed on the top of ANARCI (Dunbar and Deane, 2016), and the numbered sequence is displayed in the “Sequence numbering” tab panel, with the download of the numbered sequence accessible. User can hover on the residue in the figure to show the numbering information, including residue name, IMGT numbering and the position of the residue (V or C region, and the fragment where this residue belonging to, such as framework or CDR loops). User can switch between heavy and light chain sequences by selecting options under “Choose the sequence to display”.

##### S1.1.2 Species and isotype

We annotate the most likely species and isotype for each VCAb structure as described in section S3.1.2. This information together with the mapped allele(s) are listed when users query entries in VCAb. Alternative chain types (i.e. isotype and light chain type) would be listed in a separate column if they have the same BLAST identity with the top hit.

##### S1.1.3 Antibody structure metadata

The VCAb webserver provides tabulated details of experimental methods and resolution (downloaded from PDBe API), structural coverage (Fab or full-antibody), H-L interface, disulphide bonds, geometric angles (CH1-CL angles and elbow angles, section S3.1.4, S3.1.5), and whether the structural entry contains antigen. Hyperlinks for the corresponding PDB or SAbDab webpages are also included for further inspection.

##### S1.1.4 Antibody three-dimensional structures

Three-dimensional structural viewer is integrated in VCAb by using NGLVieweR (version 1.3.2) in R. User can rotate, zoom in/out the antibody structures in the three-dimensional structure viewer. By default, only one heavy/light chain pairs would be presented. Other chains can be presented by clicking the “view all the chains with the same PDB ID” button, with the antigen chain (if any) colored in black (section S3.1.1) and other ligand(s)(colored cyan).

#### S1.2 Analysis of CH1-CL interface

##### S1.2.1 Visualization of CH1-CL interface residues

VCAb supports interactively investigation of interface residues by two approaches. By selecting a region of interest on the domain contact map of Fab region calculated for each antibody structure, interface residues in the selected box can be automatically zoomed in, highlighted in the structural viewer and allows for a closer inspect (**Figure S1**). User can also select individual interface residues tabulated in the ” H-L interface residues” tab to achieve the same effect.

##### S1.2.2 Search structures by CH1-CL interface similarity

Taking advantage of the the CH1-CL interface analysis we performed over all VCAb structures (section S3.1.4), users can search VCAb according to CH1-CL interface similarity. In ”CH1-CL Interface” panel, users can input the PDB code and chain IDs of their antibody structure of interest. VCAb entries will then

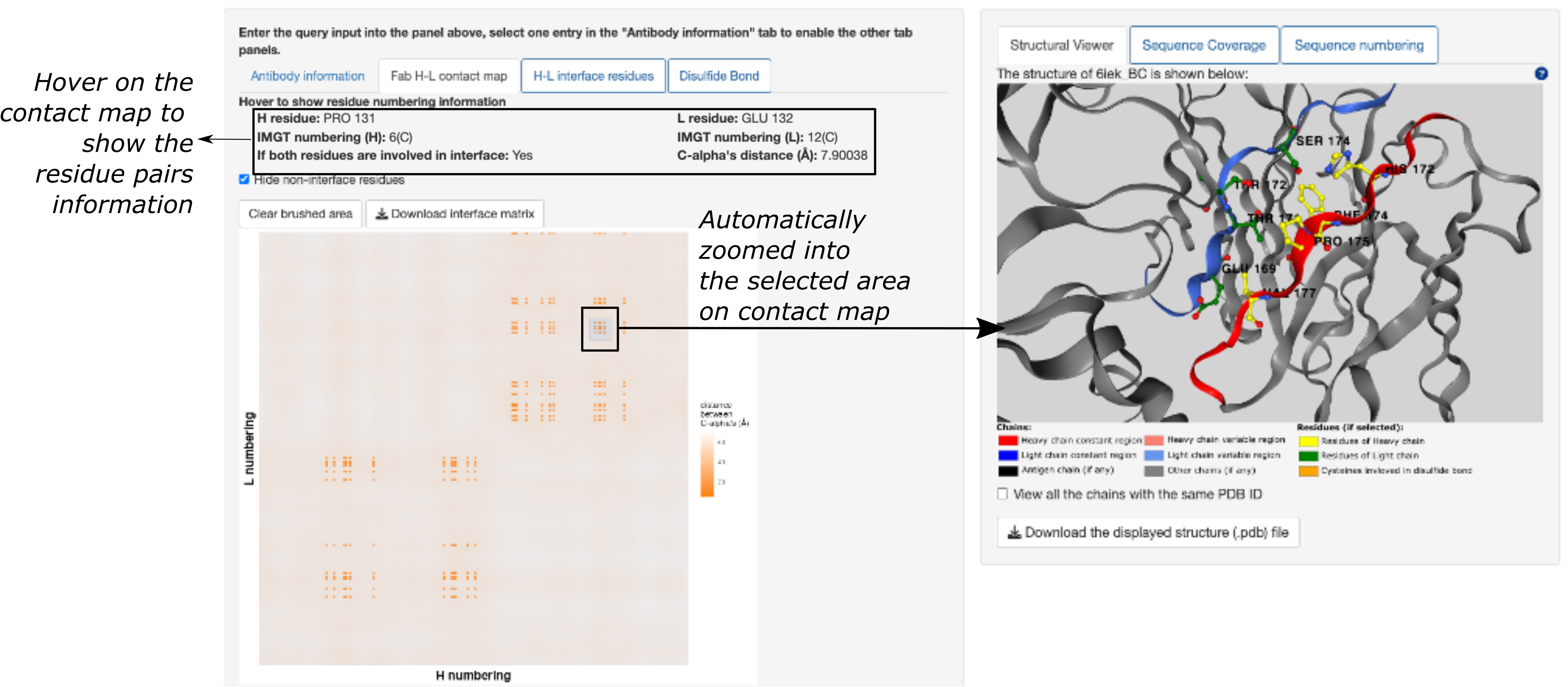

*Figure S1: Interactive domain contact map and three-dimensional structural viewer. For each VCAb structure, both the Fab H-L contact map and the three dimansional structure are displayed. The x and y axis in Fab H-L contact map represent heavy and light chain residues numbered conforming to IMGT scheme, respectively. Users can highlight residue pairs where both of them are involved in the interface by clicking ”hide non-interface residues” and show the residue information by hovering onto the contact map. The 3D structural viewer will be automatically zoomed into the selected area on the Fab contact map.*

be ranked by calculated IntDiff score (equation 4), with the smaller value indicating more similar CH1-CL interface.

#### S1.3 Search structures by antibody sequence similarity

VCAb web tool enables the filtering of antibody structures by sequence similarity, using BLAST (Altschul *et al.*, 1990) provided in the rBLAST package (version 0.99.2). This search is flexible to the region of interest (V region or both V & C regions) and both paired and unpaired H/L chain sequences. As described in section S3.1.2, sequences for V regions are extracted from the BLAST alignments and collected. Together with the sequences of the VCAb entries covering both V and C regions, four databases are formed, containing the sequences of all VCAb entries covering (i) VH domains, (ii) VL domains, (iii) full-length H chains and (iv) L chains, respectively. The webserver automatically chooses the database for the purpose of BLAST, according to the region of interest and the searching mode (paired or unpaired H and L chains) selected by the user (**Table S1**).

*Table S1: The selection of database for BLAST implemented in the VCAb application. “VH or VL” means selecting VH or VL VCAb sequences as the BLAST database, according to the chain type (H or L) selected/labeled by user. “VH and VL” means the database combining VH and VL sequences.*

**Unpaired**

**Paired**

*Fullsequence*

*Vregion*

*V&C*

*)*

*(*

*Fullsequence*

*Vregion*

*(*

*V&C*

*)*

**Searchindividualsequence**

VHorVL

HorL

HorL

VHorVL

**Searchinbatch**

VHandVL

HandL

HorL

VHorVL

VCAb supports input of single sequence and batch uploads of sequences. If individual sequence is provided, VCAb entries will be ranked by sequence identity, and the IMGT numbering for the user-inputted sequence would also be displayed. If individual sequence of V region is inputted, the C region of selected structural hits can be appended for the purpose of antibody design and/or lab investigation, with the appended sequence numbered under IMGT-scheme and downloadable.

VCAb webserver enables batch searching of sequence similarity, by allowing the user to upload a FASTA file containing multiple sequences (maximum 200 per batch). For each query sequence in the uploaded file, the top three VCAb entries given by BLAST would be listed. Under the paired mode, users need to indicate the pairing between individual H and L chains by following the format to specify the FASTA header of the sequences: AbName-HorL, where AbName should be the name of the antibody, and HorL should be either letter “H” or “L”to specify H or L chain. Paired chains can contain only one H and one L chain. Unpaired sequences would not be annotated; they are tabulated below the results of searching the paired chains.

#### S1.4 Map repertoire sequence to experimental structure

Currently, VCAb supports the upload of repertoire files and searching of matching experimentally determined antibody structures based on sequence homology. Uploaded repertoire data can be standard (AIRR, 10X Genomics CellRanger) or customised formats. When repertoire data in customised format is provided, columns holding the amino acid sequence, identity of the chain (heavy or light chain), and cell barcode should be specified, while these columns are automatically selected when standard repertoire formats are uploaded (**Table S2**). Repertoire file can be uploaded to VCAb through multiple ways: a) browsing for a local repertoire file and upload it to the server, b) live imports from online analyses on the BRepertoire (Margreitter *et al.*, 2018) webserver, or c) uploading a remote repertoire file by providing a http address. User can select different mode in searching: unpaired (individual sequence selected from sampled repertoire) or paired mode (paired heavy and light chains of the same antibody). In the unpaired mode, the sequence entry selected will be inputted into the BLAST(Altschul *et al.*, 1990) pipeline, while the latter mode automatically picked up the other paired H/L chain indicated by the same cell barcode, and input the paired chains into BLAST process.

*Table S2: Columns required for repertoire search for structural match*

**Columns**

**AIRR**

**10**

**XGenomics**

**CellRanger**

**customisedformats**

| **Amino acid sequence** | automatically selected *Column:*  *sequence alignment aa* | automatically selected *Columns: fwr1, cdr1, fwr2, cdr2, fwr3, cdr3, fwr4*; sequence fragments in these columns are automatically concatenated together for search | Multiple columns can be selected here, sequence fragments in these columns will be concatenated according to the input order |
| --- | --- | --- | --- |
| **Identity of the chain** | automatically selected  *Column: locus* | automatically selected  *Column: chain* | column values can only contain ”IGH”, ”IGK”, or ”IGL” |
| **Cell ID/ Cell barcode** ** this column is only required under paired searching mode* | automatically selected  *Column: cell id* | automatically selected  *Column: barcode* | column values contain unique cell ID for each  cell |

Structural hits will be ranked by percentage identity of the amino acid sequence to the selected individual or paired chains in uploaded repertoire. Antibody and structural information for each structural hits will be displayed, with its three-dimensional structure visualised and inspected (section S1.1).

As an example, we showed structure matches of the two sequences we inputted from a published antibody sequence repertoire (Stewart *et al.*, 2022):

- The top structural match (PDB 7tn0) of the repertoire sequence with identifier

“CV222D2 M 10050314 TAACCTTCTGTTATAT RD39” indicates antibody recognise RBD epitope distinct from ACE2 binding site.

- Another sequence with identifier “CV227D9 G1 10089371 TTCGGTCACGTGAATT RD30” is also inputted to VCAb to search for structural match (PDB 7bz5), which shows this antibody directly blocks ACE2 binding.

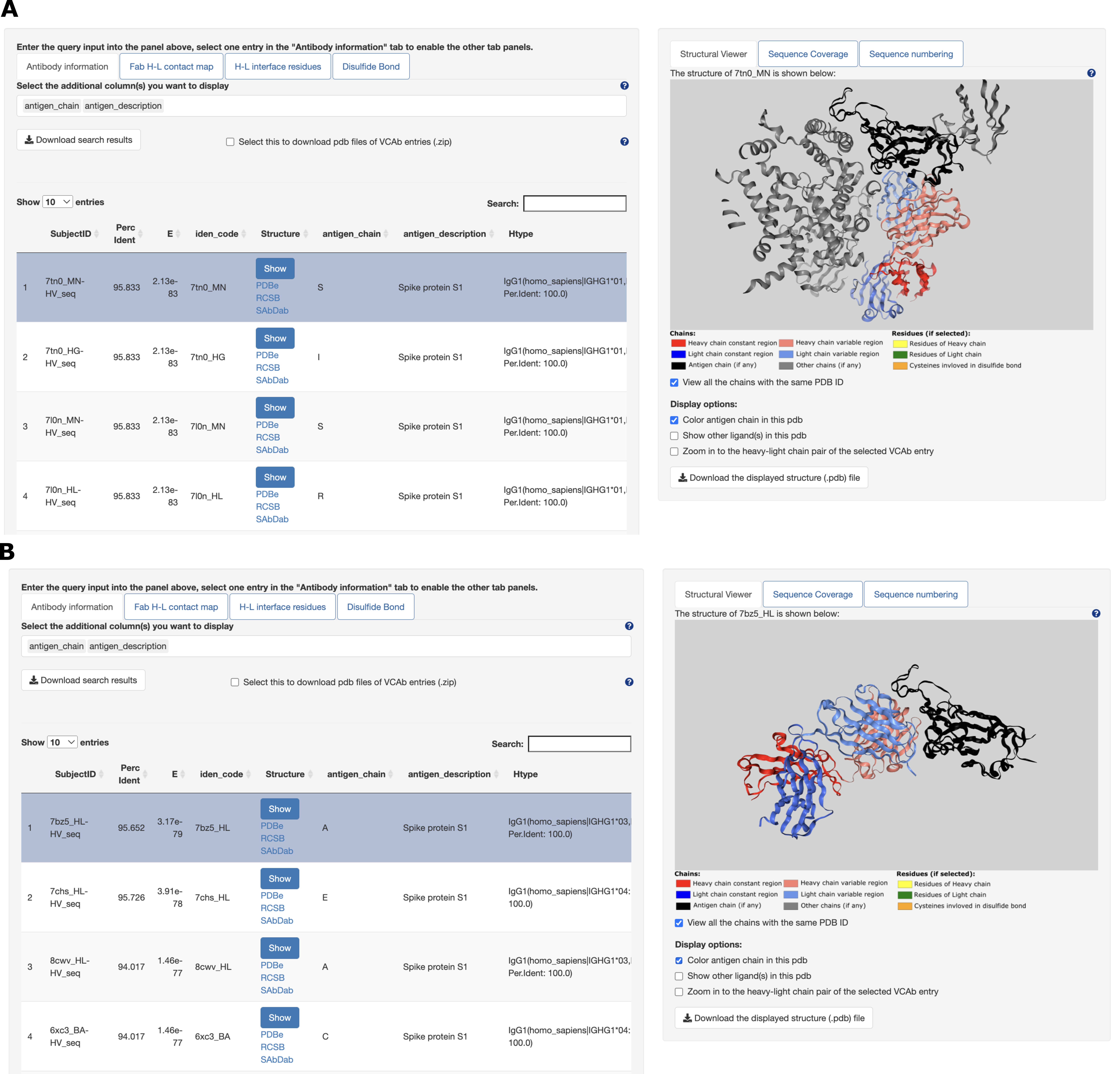

*Figure S2: Investigation of COVID-19 repertoire from a structural perspective. (A) Top structural match (PDB 7tn0) of the repertoire sequence “CV222D2 M 10050314 TAACCTTCTGTTATAT RD39” indicates antibody recognise RBD epitope distinct from ACE2 binding site. (B) Repertoire sequence*

*“CV227D9 G1 10089371 TTCGGTCACGTGAATT RD30” is inputted to VCAb to search for structural match (PDB 7bz5). The top structural match shows this antibody directly blocks ACE2 binding.*

#### S1.5 Access pre-computed *in silico* mutational scanning data

##### S1.5.1 Visualization of the *in silico* mutational scanning results

For each structural hit, pre-computed *in silico* mutational scanning profiles for the heavy and light chains have been generated by Rosetta pmut scan, AntiBERTy, and AlphaMissense. For each wild-type (WT) residue on the sequence, its mutation to every one of the other 19 amino acids has been evaluated using these methods to yield a score measuring mutational impact. These data are visualised as heatmaps which are displayed in separate tabs. Each *in silico* mutational scanning heatmap is accompanied by a residue selection strip positioned above the heatmap (**Figure S3**). Each dot on the residue strip represents one amino acids on the displayed structure, corresponding to one column in the heatmap. Residue selection strip is designed to allow the user to select residues of their interest to highlight on structural viewer. CDRs/loops and frameworks are highlighted by alternating dark/light colours on the selection strip. The x-axis for the heatmap represents the antibody sequence, and the y-axis represent the 20 amino acids. Red pixels in the heatmap represents mutations which are not preferable, and blue pixels represent preferable mutations. The

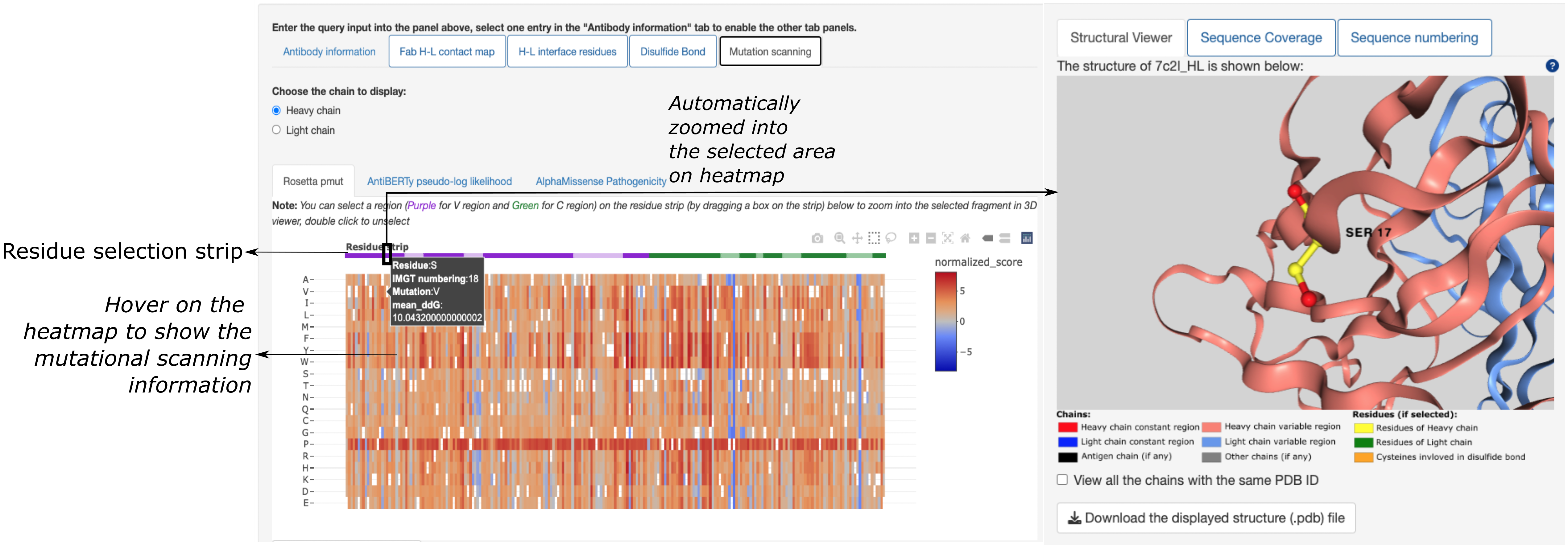

*Figure S3: in silico mutational scanning profiles and three-dimensional structural viewer.*

WT residue for each position is marked by white. A hovered box is displayed for each point in the heatmap, showing the WT and mutant, the IMGT numbering of the position and the score representing the mutational impact as evaluated using the method selected by the user.

##### S1.5.2 AntiBERTy mutation score

AntiBERTy mutational scanning score is derived from the pseudo-log likelihood of each possible amino acids being presented at every position, as determined by the AntiBERTy language model trained over 26 million antibody sequences (Ruffolo *et al.*, 2021). We scaled the raw AntiBERTy score by subtracting the AntiBERTy score for the wild type (WT) residue at each position, such that they can be interpreted as relative changes in amino acid preference compared to WT (equation 1). Since these likelihood scores are in the logarithm space, the mutation score of multiple co-existing point mutations is calculated by summing up scores corresponding to the individual mutations (equation 2, **Figure S4**).

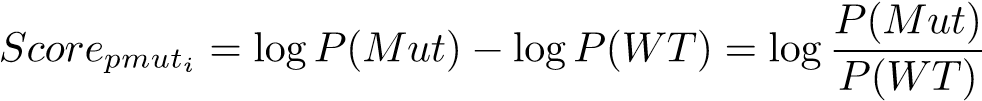
(1)

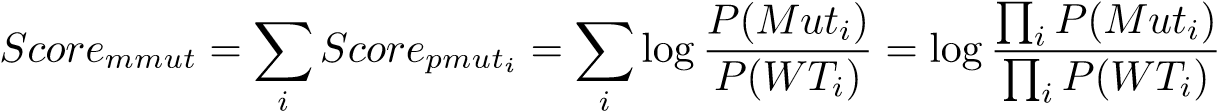
(2)

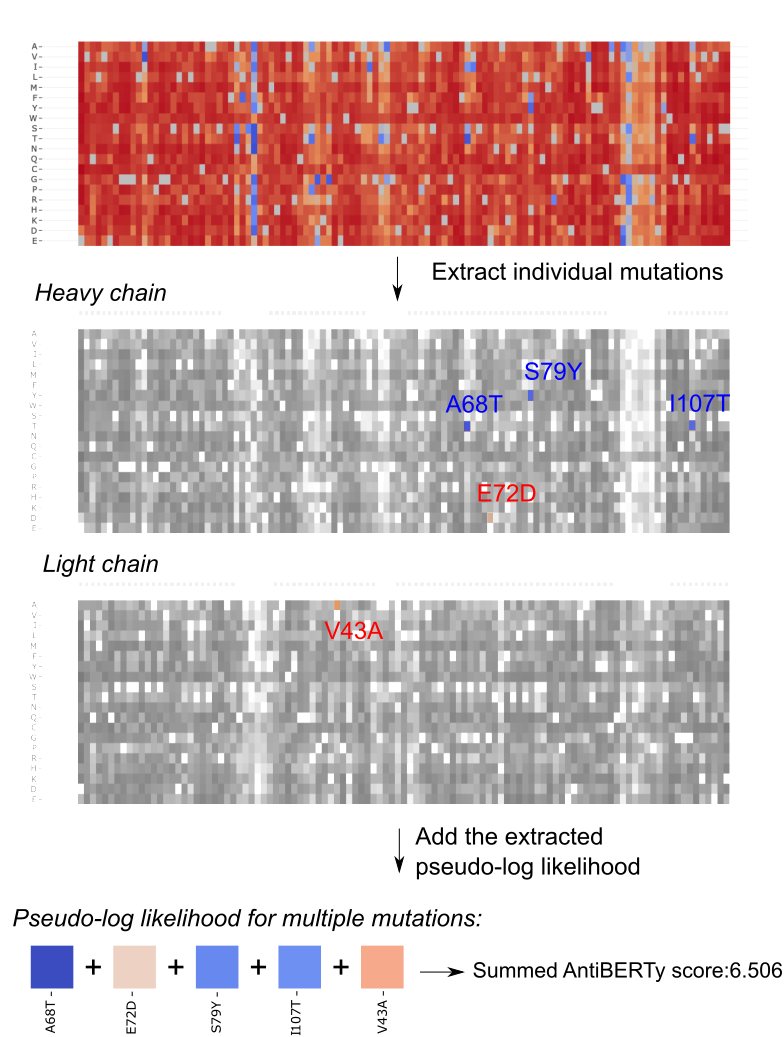

*Figure S4: AntiBERTy mutational score can be added together to represent the mutational impact of multiple co-existing amino acid substitutions.*

### S2 Case studies

#### S2.1 Interface Investigation

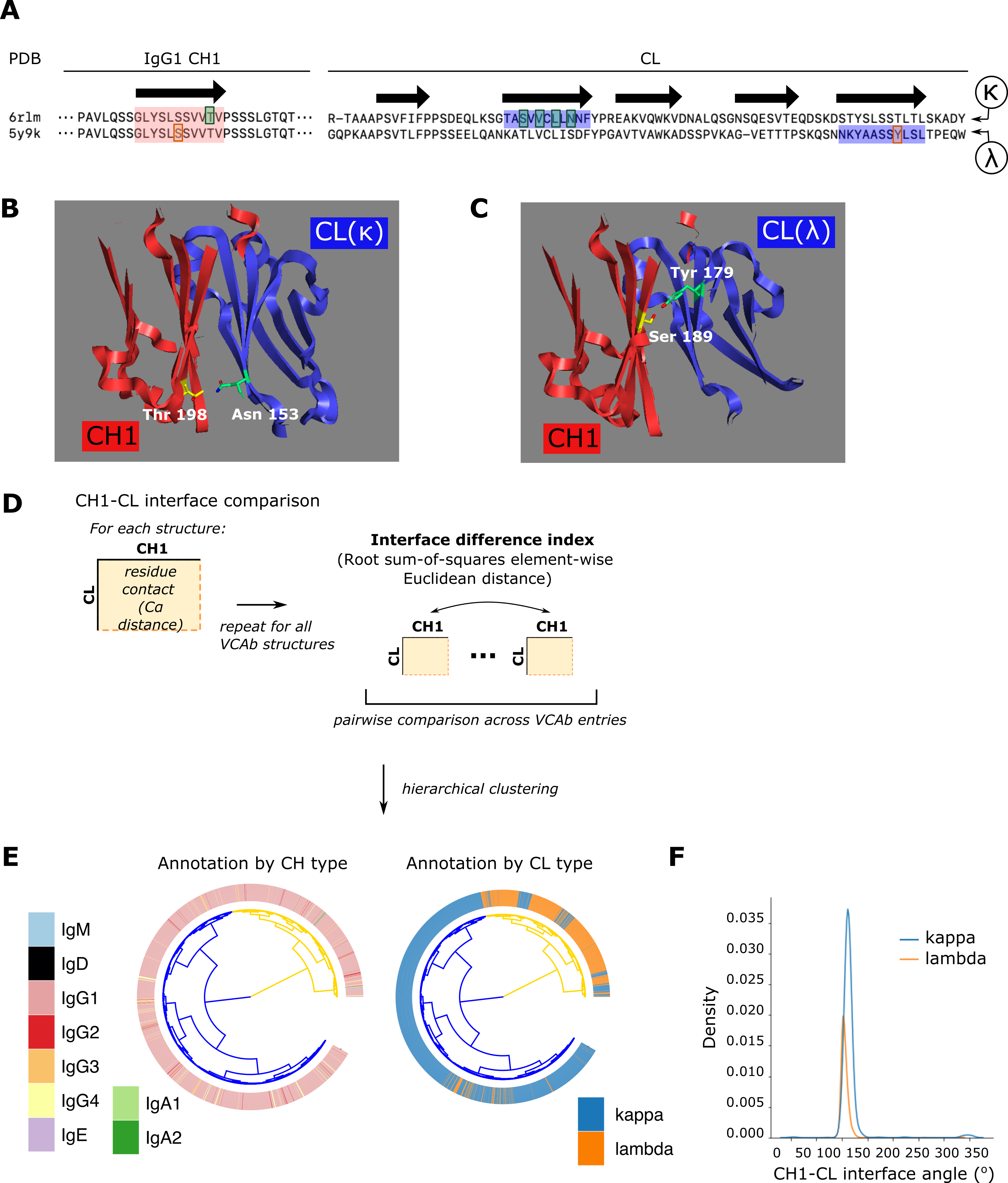

*Figure S5: CH1-Cl interface analysis. (A) The sequence alignment for the CH1 and CL sequences of PDB 6rlm (IgG1-κ) and PDB 5y9k (IgG1-λ). Arrows at the top of the sequence represents the position of the beta strand in the domain. Regions involved in the CH1-CL interface are highlighted in red (CH1) and blue (CL), with specific interface residues indicated with dark coloured boxes. (B and C) The display of the interacting residues of interest between CH1 and CL(κ) from PDB 6rlm (B), and CH1 and CL(λ) from PDB 5y9k (C). (D) Schematic explaining the process to compare CH1-CL interface across VCAb entries, by comparing residue contacts between CH1 and CL to generate what we term the “interface difference index” which compares CH1-CL Cα contacts between any pair of VCAb structures. (E) Hierarchical clustering analysis of pairwise interface difference index across all human VCAb entries reveals 2 clusters of CH1-CL residue contact patterns. Each structure is annotated by the CH (left) and CL (right) types. (F) Distribution of CH1-CL interface angle for VCAb entries using the κ (blue) and λ (orange) light chains.*

Analysis of the CH1-CL interface is important in our understanding of the principles behind the pairing between H and L chains, and the design of bispecific antibodies for therapeutic purposes (Lewis *et al.*, 2014; Liu *et al.*, 2015; Mazor *et al.*, 2015; Krah *et al.*, 2017). With VCAb encompassing accurate annotations of the C region of antibody structures, we hypothesise that the database can be used to compare the residue composition of the CH1-CL interface under different types of H-L combination. Here, we give an example of comparing CH1-CL interface residues when different types of light chain (*κ* or *λ*) is paired with the same type of heavy chain (IgG1). The PDB entry 6rlm is an IgG1-*κ* antibody, while PDB 5y9k is IgG1-*λ* antibody. As expected, the CH1 sequences represented in these two structures are identical (**Figure S5A**). However, our analysis suggests that in these structures the *κ* and *λ* chains use different *β*-strands in engaging with the IgG1 H chain, involving different residues: in the IgG1-*κ* structure Thr198 of the heavy chain is in the interface with the CL domain (**Figure S5B**), while this threonine residue in the IgG1-*λ* structure is not (**Figure S5A**). We observe the opposite scenario for a nearby serine residue at the same *β*-strand: Ser189 in the IgG1-*λ* structure is involved in the CH1-CL interface (**Figure S5C**) while the serine residue at the same position in the IgG1-*κ* structure is not (**Figure S5A**). This indicates that in these two cases, the heavy chain of the same sequence in IgG1 pairs with the *κ* and *λ* chains using a slightly different configuration, with a certain degree of compensation from amino acids with similar physicochemical characteristics in the vicinity of one another.

Beyond this single example, VCAb provides the opportunity to investigate such problems at a larger scale with its accurate annotation of such structural features, periodically updated to include newly resolved antibody structures. We compare the residue contact patterns at the CH1-CL interface (**Figure S5D&E**) across all human VCAb entries. We performed pairwise comparison of the residue contacts between any two Fab structures (formula defined in **equation 4**) to generate a distance matrix. Hierarchical clustering (using

Ward’s distance in R) was applied to this distance matrix to generate the dendrogram shown in **Figure S5E**, using the ggtree package. This comparison reveals two CH1-CL residue contact patterns; we found that whilst these patterns do not seem to depend on heavy chain type (primarily because of the sparsity of structures other than those of the IgG1 subtype in the data), the interface residue contact patterns can be separated in accordance with the light chain types of antibody: most antibodies with *κ* are within one cluster, while most antibodies with *λ* are clustered together (**Figure S5E**), suggesting the choice of light chain partner influences CH1-CL packing. We further investigate the geometry of this CH1-CL packing by comparing the CH1-CL interface angles for antibodies with *κ* and *λ* light chains (**Figure S5F**). The difference between *κ* and *λ* paired antibodies is small (median for *κ* structures ≈ 112^◦^, median for *λ* ≈ 104^◦^). We also considered the elbow angles and it displays a similar pattern (**Figure S5F**). In conclusion, CH1-CL interface patterns appear to be influenced by the light chain, utilising different residue contacts for such packing without altering the packing angles governing Fab geometry. In the VCAb web-server, users can query an antibody structure to search for other structures with similar CH1-CL residue contacts; these structures can again be interactively visualised and downloaded via the webserver.

### S3 Methods

#### S3.1 Data collection and annotation

##### S3.1.1 Collecting Antibodies

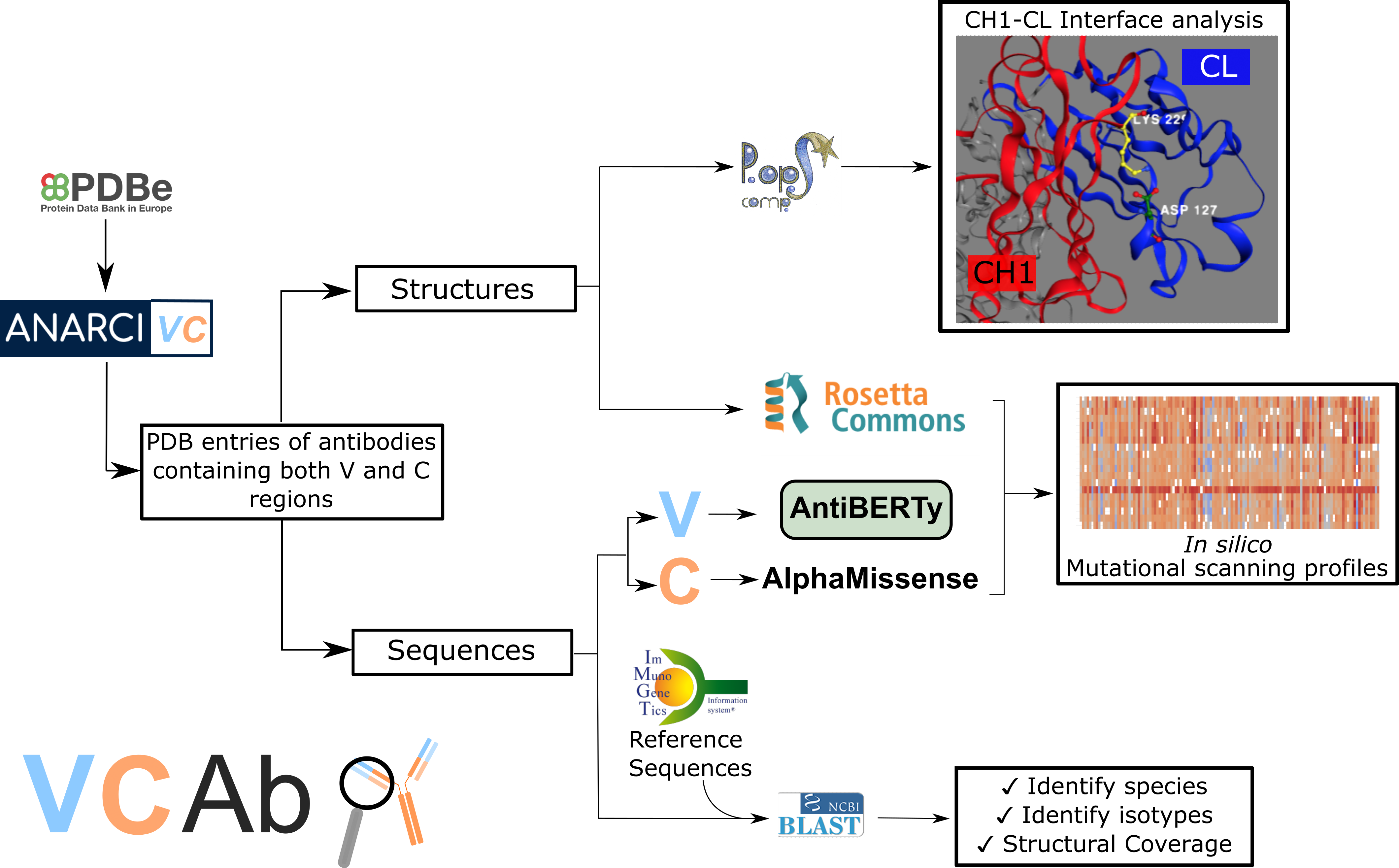

*Figure S6: Schematic of the data collection, data annotation, and structure analysis in VCAb.*

Protein sequences are downloaded (download date: 18th March, 2024) from worldwide PDB archive(Burley *et al.*, 2019) and inputted into ANARCIvc (section S3.1.3), a package modified from ANARCI (Dunbar and Deane, 2016), to identify all the antibody sequences containing both V and C regions in PDB (**Figure S6**). Heavy and light chains in each PDB entry are paired by applying the distance constraint of 22 ˚A between the conserved cysteines at position 104 of IMGT antibody V region numbering (Dunbar *et al.*, 2014; Lefranc *et al.*, 2005). Antigen chains are recorded as the chain bearing fragments within 7.5 ˚A to the CDR loops of the antibody (Dunbar *et al.*, 2014).

We note that the sequence directly accessed from PDB archive does not necessarily match the sequence represented in the PDB ATOM records and this could be misleading when selecting template for modelling. We therefore downloaded the structural files (mmCIF format) and extracted the ATOM records for accurate feature annotation. Residues missing ATOM records are omitted in this case. Sequences downloaded directly from PDB archive and sequences extracted from the ATOM records are used for different purpose (see section

S3.1.2).

To remove redundancy due to multiple copies of the same antibody sequence being represented in the PDB structures, antibody entries are collapsed based on whether they have the same PDB identifier and the same coordinate sequences (i.e. the sequence represented in the PDB ATOM records) of both H and L chains. We give a unique identification code for each antibody entry, in the format of pdb HL, where pdb is the 4-character PDB code, H and L are the chain IDs of respective heavy and light chain from the first H-L pair. Duplicated antibody chains found in the same assembly are recorded in the collapsed database separately with all matching chain names.

##### S3.1.2 Species, isotype and structural coverage annotation

We noticed that species annotation for antibody sequences can be highly complex, and in some cases the single species nominated in the annotations provided by the PDB can be misleading. For example, some antibodies annotated as human in PDB might contain sequence fragments from other species, such as chimeric antibodies (i.e. sequences of V and C regions are from different species) or humanised antibodies (i.e. antigen-binding loops and fragments are grafted onto a human immunoglobulin framework). These antibodies are reflected by the low percentage identity when aligning to the human reference sequences for either/both of the C and

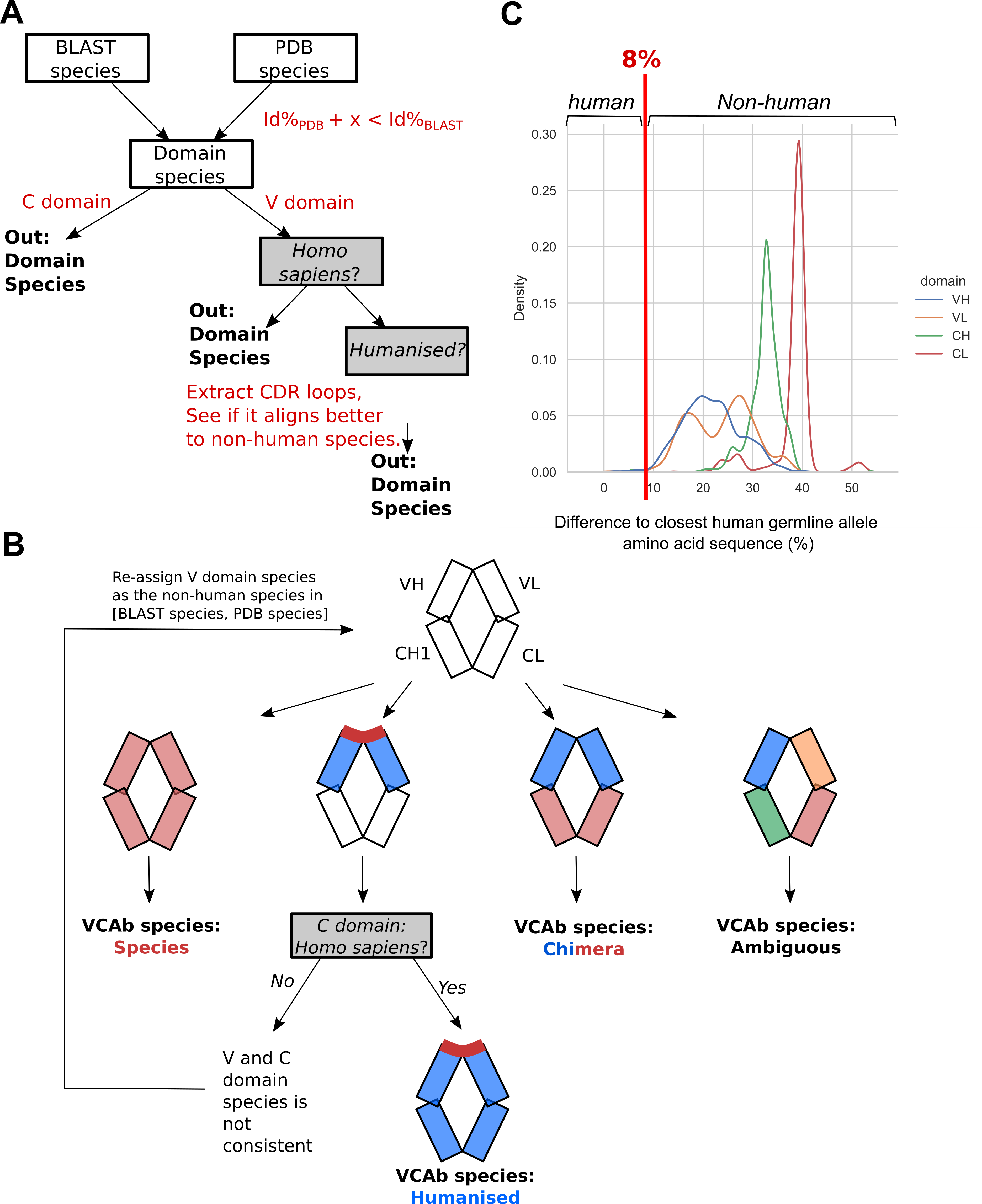

*Figure S7: Species assignment in VCAb. (A) The process to assign species to domains. BLAST species is the species which give the highest percentage identity when aligned to the reference gene, PDB species is the species annotation in Protein Data Bank. We compare the percentage identity of the top hit from the BLAST species*

*(Id*%*_BLAST_) to that of the top hit from the PDB species (Id*%*_PDB_). To allow for mutations and sequence polymorphisms, we only overwrite the PDB species as the designated species of the domain if the percentage identity for the BLAST species is higher than a certain value (x) compared with the Id*%*_PDB_. For V domains, an additional step is applied to identify whether it is humanised, bu extracting CDR loops (section S3.1.2). (B) Procedure to assign the species for the entire antibody structure. Species are assigned to each domain first by the process shown in panel A, then all the domains species are considered together to give the species for the antibody. Different colors indicate different domain species, the dark red lines indicate the CDR. (C) The distributions for non-human antibodies of the difference between Id*%*_BLAST_ and the Id% when aligning to human reference genes. These sequences are from non-human antibodies where PDB species and BLAST species are consistent, and is trying to mimic the situation when they are mistakenly annotated as human. Different colors show the distribution of different domains. Majority shows the distribution higher than 8%. We therefore set x as illustrated in panel A as 8 in our pipeline.*

V regions. To flag these cases, sequences of V and C regions for both H and L chains are extracted and passed as input to separate BLAST runs, to compare with the reference alleles from different species of V and C regions (downloaded from IMGT, date accessed: 1st July, 2022), and search for the alleles with the best sequence similarity. Species is firstly assigned for V and C regions separately (**Figure S7a**), then the final decision of the species for the entire antibody is made from the species annotation for regions (**Figure S7B**).

By doing so, we flag chimeric antibodies as those with different species annotations for the V and C regions. If an antibody is identified as human, then the CDR loops (IMGT definitions) would be extracted to compare its alignments to reference alleles from other species versus human, to identify humanised antibodies (**Figure S7ac**). Species annotation provided by PDB will be overwritten by the species of the best BLAST hit when the percentage identity for the best BLAST hit is higher than the hit corresponding to the PDB species by a certain cut-off (**Figure S7A**). To determine this cut-off, we extracted the non-human antibodies where the species annotations derived from BLAST and PDB are consistent. Sequences of non-human antibodies are aligned with the human reference sequences, to mimic the situation if the non-human antibodies are annotated as human in PDB. Then the percentage identity (Id%) for species from the best BLAST hit are compared with the Id% when aligned to human references. We find that a cut-off of 8% effectively distinguishes non-human and human antibodies (**Figure S7C**).

To identify the isotype of antibody chains found in the collated VCAb entries, we search sequences directly derived from PDB SEQRES records against IMGT (Lefranc *et al.*, 2015) reference CH and CL alleles of the species we assigned based on the procedure detailed above, using the BLAST program in the command line via Bio.Blast.Applications module in Biopython (version 1.79) (**Figure S6**). Alleles of isotypes downloaded from IMGT are first collapsed by their amino acid sequences to build a non-redundant BLAST sequence database. The best BLAST alignment with highest percentage identity is considered to assign the CH/CL type of the query chain.

The isotype of the antibody is listed in the Htype column of the VCAb database. In this column, following the name of isotype, the allele with the highest percentage identity in the BLAST result is shown. Alleles with identical amino acid sequence are also listed. For cases where the sequence is similarly close to multiple isotypes to give the same percentage identity or score as joint top hits in BLAST, they are listed in the Alternative Htype column for users’ reference. The light chain type of antibody is identified in the same process likewise.

To identify the structural coverage of the antibody, the heavy chain sequence represented in PDB ATOM records (instead of the sequence from PDB SEQRES records) are searched using BLAST against the IMGT reference IGHC sequences for the assigned C region species (**Figure S6**). Based on the start and end positions of the BLAST alignments, together with the domain boundaries described in the FASTA header of relevant CH alleles in IMGT, we classify the structural coverage of the antibody as Fab (the heavy chain contains only VH and CH1, but no other domains) or full antibody (the heavy chain contains every CH domain relevant for the given isotype). Since the light chain is immaterial in the classification into Fab and full antibodies (in both cases VL and CL should be present), this classification is based on the domain coverage of the heavy chain sequence with ATOM records extracted from PDB files. A second check is applied here to make sure all the antibodies in VCAb contain both V and C regions, by excluding antibodies identified only containing V regions using the sequence observed in PDB ATOM records; users can download the list of structures which are excluded from VCAb due to failures to meet these listed criteria on the VCAb web application.

The sequences of antibodies are separated into V and C regions, based on identifying the beginning of the alignment to the CH/CL alleles. These sequences for V region are collected into a flat-file which can be used as a sequence database for searching VCAb based on the V sequence similarity (see section S3.3, “VCAb webserver”).

##### S3.1.3 Antibody C region numbering

Antibody sequence numbering is a process to assign, for every observed residue in the antibody sequence, an identifier based on a scheme that can be applied to every antibody sequence. The use of such universal scheme would allow cross-comparisons of residues between structures, for convenient extraction of its structural position (e.g., with respect to secondary structure elements, or for structurally conserved positions such as cysteines which precede CDR loops) and comparisons of these features across different antibody sequences. Numbering schemes for the V region have been well-established with an array of computational tools already available that automate the process to number a user-supplied V region sequence. Although a numbering scheme is also curated for the C region (IMGT)(Lefranc *et al.*, 2005), to our knowledge there are no tools which allow users to automatically number C region antibody sequence. We reason this is essential to generate annotations in VCAb which can be interpreted for sequence characterisation: to identify antibodies with C region sequences (see section S3.1.1, “Collecting antibodies”), to define reference points for CH1-CL angle calculation, “CH1-CL interface angle”), and to compare CH1-CL interfaces across VCAb (see section

S3.1.4).

We address this issue by modifying the tool ANARCI (Dunbar and Deane, 2016). ANARCI built Hidden Markov Models (HMMs) of V region germline sequences (Dunbar and Deane, 2016), which can subsequently be used to number a user-supplied V region sequence using the hmmscan function in HMMER (Dunbar and Deane, 2016). We expand ANARCI to ANARCIvc to enable the numbering of both V and C region sequences. For the numbering of C sequences, similarly to ANARCI, a set of HMM profiles are generated for reference sequences of CH1 and CL domains from different species automatically downloaded from IMGT. The source code of ANARCI is modified in order to implement the IMGT numbering scheme for the C region, and apply this scheme to number the C region observed in any antibody structure during the VCAb compilation process. The IMGT C region numbering scheme is built on the basis of V region numbering scheme, where it inherits the V region numbering for structurally equivalent *β* strands when structures for V and C domains are superimposed, remove the numbering for extra beta strands in V domains, and place insertions for additional residues in C domains(Lefranc *et al.*, 2005). The implementation and HMM models for V region numbering has been wholly inherited from ANARCI without modification. ANARCIvc source code is freely available as a github repository forked from ANARCI: [https://github.com/Fraternalilab/ANARCI_vc.](https://github.com/Fraternalilab/ANARCI_vc)

##### S3.1.4 CH1-CL interface analysis

**H-L interface residues** POPSComp (Kleinjung and Fraternali, 2005) is applied to identify the residues involved in the interface between heavy and light chains, by evaluating the difference in the solvent accessible surface area (SASA) upon the formation of the H-L complex (**Figure S6**). This SASA difference (∆SASA) is the area buried due to the interaction between the H and L chains. Residues with ∆SASA larger than 15 ˚A^2^ is considered as part of the H-L interface and listed as a table in the VCAb webserver; this ∆SASA cut-off has previously been applied to study protein-protein interaction interfaces across the PDB (Fornili *et al.*, 2013).

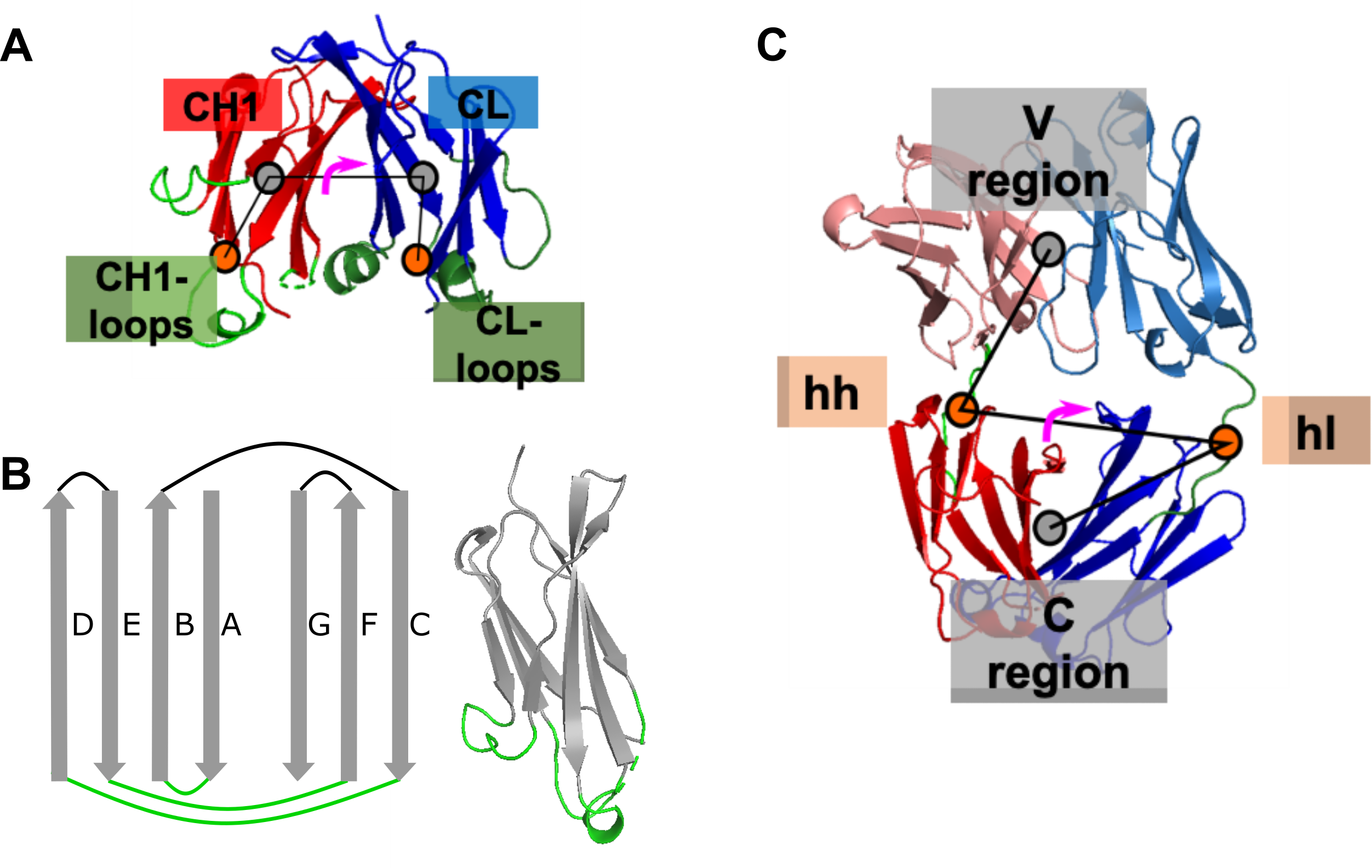

*Figure S8: Angles calculated in VCAb. (A) CH1-CL interface angle defined by then center of mass (COM) of CL loops, CL domain, CH1 domain and CH1 loops. (B) CH1 and CL loops is defined as the loops between β-strand A&B, C&D, E&F for both CH1 and CL, and is highlighted in green. (C) The elbow angle is calculated as the torsion angle of the COM of V region, the COM of the loop linking VH and CH1 domain (hh), COM of the loop linking VL and CL domain (hl), and COM of the C region of the Fab.*

**CH1-CL interface angle calculation** CH1-CL interface angles is defined as the torsion angle formed by the center of mass (COM) of the CL loops, COM of the CL domain, COM of the CH1 domain, and COM of the CH1 loops (Fern´andez-Quintero *et al.*, 2020) (**Figure S8A**). The CH1 and CL domains are extracted based on the domain boundaries identified during the BLAST step (see section S3.1.2). The region of CH1 loops and CL loops are defined as the fragments linking adjacent *β*-strands at the C-terminal of the C domain, i.e. fragments between *β*-strand A&B, C&D, E&F for both CH1 and CL (**Figure S8B**, (Chiu *et al.*, 2019)). The extraction of loops are based on loop positions suggested by antibody C region numbering for each VCAb entry. After these, the COM for CL loops, CL domain, CH1 domain and CH1 loops are calculated using Bio.PDB package (Bio version 1.79), for the calculation of CH1-CL angle.

**CH1-CL interface similarity comparison** To compare patterns of residue contacts between the CH1 and CL domains across all VCAb entries, we first generate a matrix *r* storing pairwise C*α* distances between residues in the CH1 and CL domains; in the formula below, *h* and *l* refer to residues from the CH1 and CL domains, respectively.

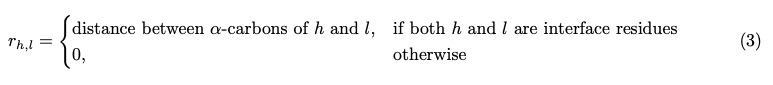

Interface residues are taken from POPSComp analysis. In order to keep the dimensions of every contact matrix constant across all VCAb entries, sequences of CH1 and CL domains for all VCAb entries are numbered by ANARCIvc, and the resultant IMGT-gapped sequences are used for the two dimensions of the residue contact matrix.

We then compute a distance metric to quantify the difference between any two given residue contact matrices from two VCAb entries. The “interface difference index” (IntDiff(*A,B*)) is calculated for every pair of VCAb entries using the formula below, where A, B are residue contact matrices of VCAb entries:

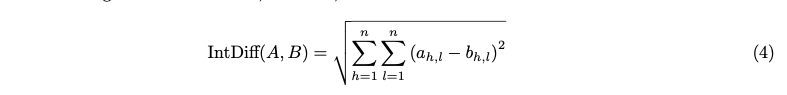

This amounts to a root sum-of-squares element-wise difference between two matrices *A* and *B*, which is used to search for antibody structures with similar CH1-CL contact in VCAb web server, or potentially for large-scale analysis (section S2.1).

##### S3.1.5 Elbow angle calculation

Elbow angle is defined as the torsion angle set by the center of mass (COM) of the V region (VH and VL domains), COM of the loop linking VH and CH1 domain, COM of the loop linking VL and CL domain, and COM of the C region of the Fab (CH1 and CL domains)(Fern´andez-Quintero *et al.*, 2020) (**Figure S8C**). V region and C region of the Fab are extracted according to the domain boundaries identified in BLAST (see section S3.1.2). The boundaries of loops linking VH and CH1, VL and CL are set by finding the nearest residues involved in adjacent *β*-strands with respective to the VH-CH1, VL-CL boundaries identified in BLAST. The extraction of these regions and the calculation of COM are performed using Bio.PDB package (Bio version 1.79).

##### S3.1.6 Disulphide bond calculation

The distance between C*α* of cysteines involved in disulphide bond is below 7.5 ˚A in most cases (Gao *et al.*, 2020). Distances between C*α*s of all the cysteines from each VCAb entry are calculated, and 7.5 ˚A is used as threshold to consider cysteines as forming a disulphide bridge.

#### S3.2 Isotype sensitivity analysis using *in silico* mutational scanning data

Rosetta pmut scan results can be downloaded from the VCAb webserver, for further analysis of positions in VH domain where the stability changes significantly upon isotype change. We reason that these positions potentially represent regions sensitive to switching the CH1 domains from one isotype to another. Isotype switching is an important process for B cell maturation *in vivo*, and also essential in the design of antibodies to accomplish specific effector functions. To mimic this process *in silico*, we selected as case study two antibody structures with the same V region sequence but different isotype (IgA1: PDB 3qnx; IgG1: PDB 3qo0). We additionally selected another IgG1 with the same V region (PDB 3qo1) as a negative control. This set of structures allow us to perform two comparisons: IgA1 v.s. IgG1 and IgG1 v.s. IgG1. The goal here is to find positions that produce significant changes in IgA1-IgG1 compared with IgG1-IgG1.

Within each comparison (i.e. IgA1-IgG1 or IgG1-IgG1), the difference between the mutation scores of two structures are calculated (∆*mut*), which is used to generate the Volcano plot in the main text (Figure 4). For each wild-type residues, Student’s t-test is performed to calculate the p-values by comparing the 19 values of ∆*mut* in group IgA1-IgG1 (∆*mut_A_*_1_*_,G_*_1_) and group IgG1-IgG1 (∆*mut_G_*_1_*_,G_*_1_). The fold change (FC) for each wild-type residue (*r*) is defined as equation 5, i.e. obtaining the excess mutational impact (represented by the average mutational difference over all 19 mutations for a given residue) in the IgG1-IgA1 comparison ($\bar{{\Delta mut}_{A1,G1}}$) over that from the IgG1-IgG1 comparison ($\bar{{\Delta mut}_{G1,G1}}$).

We note further that the FC*_r_* value can be either positive or negative: since high mutation scores from Rosetta pmut represent low stability, positions with FC larger than zero indicates mutations here tends to be IgG1 favorable and FC smaller than zero indicates IgA1 favorable. We therefore normalise these FC*_r_* values by an inverse hyperbolic sine transformation (i.e. *arcsinh*(FC)). In the volcano plot (main text Figure 4C), the x-axis depicts *arcsinh*(FC) while the y-axis represents −log_10_ p-value. Each dot represents a wild-type residue; those with |FC| larger than 2 and p value smaller than 0.01 are highlighted.

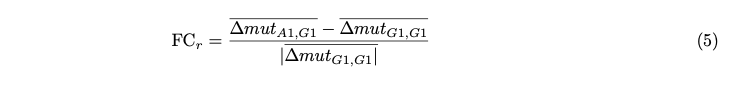

#### S3.3 VCAb webserver

The website version of the VCAb is built to allow public access for academic research purposes. Shiny (version 1.6.0) in the R statistical computing environment (version 4.1.1) is used as the basis to build the website. NGLVieweR (version 1.3.2) is used for visualisation of antibody structures. The VCAb webserver is available at: [https://fraternalilab.cs.ucl.ac.uk/VCAb/.](https://fraternalilab.cs.ucl.ac.uk/VCAb/) The underlying database is updated monthly, with the automatic pipeline scheduled to start at 07:00 Greenwich Mean Time on the last Monday of every calendar month. The pipeline begins by download all the protein sequences in PDB archive, and perform all the aforementioned analyses to populate VCAb. The source code for generating and hosting the shiny VCAb webserver is also made available ([https://github.com/Fraternalilab/VCAb)](https://github.com/Fraternalilab/VCAb) for users to host a local version of the VCAb database.

The VCAb web server provides visualisation functionalities for the queried 3D structures, structural coverage, *in silico* mutational scanning profiles, antibody numbering information for both V and C regions, tabulated details of CH1-CL interface and disulphide bond, as well as bulk download of search results and the entire VCAb database in Comma Separated Value (CSV) format. For the data tables in the “Antibody information” panel, the following columns are displayed by default: iden code, Structure (button to display in the 3D structure viewer and links to external databases), Htype (assignment of CH allele), Ltype (assignment of CL allele), Structural Coverage (indication of Fab/full antibody represented in the atomic coordinates); users can customise this view by selecting other columns (such as special cases) accessible by selecting the corresponding names in the “Select the additional column(s) you want to display” box. All columns as described in the online documentation can be accessed in the downloadable CSV files.
